# Evaluation of RNA*later*™ as a field-compatible preservation method for metaproteomic analyses of bacteria-animal symbioses

**DOI:** 10.1101/2021.06.16.448770

**Authors:** Marlene Jensen, Juliane Wippler, Manuel Kleiner

## Abstract

Field studies are central to environmental microbiology and microbial ecology as they enable studies of natural microbial communities. Metaproteomics, the study of protein abundances in microbial communities, allows to study these communities ‘*in situ*’ which requires protein preservation directly in the field as protein abundance patterns can change rapidly after sampling. Ideally, a protein preservative for field deployment works rapidly and preserves the whole proteome, is stable in long-term storage, is non-hazardous and easy to transport, and is available at low cost. Although these requirements might be met by several protein preservatives, an assessment of their suitability in field conditions when targeted for metaproteomics is currently lacking. Here, we compared the protein preservation performance of flash freezing and the preservation solution RNA*later***™** using the marine gutless oligochaete *Olavius algarvensis* and its symbiotic microbes as a test case. In addition, we evaluated long-term RNA*later***™** storage after 1 day, 1 week and 4 weeks at room temperature (22-23 °C). We evaluated protein preservation using one dimensional liquid chromatography tandem mass spectrometry (1D-LC-MS/MS). We found that RNA*later***™** and flash freezing preserved proteins equally well in terms of total number of identified proteins or relative abundances of individual proteins and none of the test time points were altered compared to t0. Moreover, we did not find biases against specific taxonomic groups or proteins with particular biochemical properties. Based on our metaproteomics data and the logistical requirements for field deployment we recommend RNA*later***™** for protein preservation of field-collected samples when targeted for metaproteomcis.

**Importance:** Metaproteomics, the large-scale identification and quantification of proteins from microbial communities, provides direct insights into the phenotypes of microorganisms on the molecular level. To ensure the integrity of the metaproteomic data, samples need to be preserved immediately after sampling to avoid changes in protein abundance patterns. In laboratory set-ups samples for proteomic analyses are most commonly preserved by flash freezing; however, liquid nitrogen or dry ice is often unavailable at remote field locations due to its hazardous nature and transport restrictions. Our study shows that RNA*later***™** can serve as a low hazard, easy to transport alternative to flash freezing for field preservation of samples for metaproteomics. We show that RNA*later***™** preserves the metaproteome equally well as compared to flash freezing and protein abundance patterns remain stable during long-term storage for at least 4 weeks at room temperature.

## Introduction

Field studies are central to environmental microbiology and microbial ecology as they allow for the *in situ* study of microbial communities and their interactions with the biotic and abiotic environment. This includes the study of symbiotic interactions between microorganisms and animal or plant hosts. Many of the approaches used to study these symbioses “*in situ*” in reality require that samples are preserved in the field for later analysis in the laboratory. This includes, for example, cell counting of specific taxa using fluorescence *in situ* hybridization (FISH) methods [1–3] and various sequencing based approaches for the characterization of taxonomic community structure and functional potential such as 16S rRNA gene sequencing [4–6] and shotgun metagenomics [7–11]. Sample preservation methods for most of these approaches have been established over the last few decades [12–14]. However, for some approaches, including metaproteomics, field preservation methods have not been extensively investigated.

Metaproteomics is an umbrella term that encompasses approaches for the large-scale identification and quantification of expressed proteins in a microbial community [15, 16]. Metaproteomics can be used not only to determine the metabolism and physiology of community members but also to estimate community member abundances, interactions and carbon sources [16–18]. Over the last decade metaproteomics has led to many significant discoveries including novel insights into microbial biofilm communities collected from acid mine drainages (AMD) [19, 20], marine symbioses [11, 21, 22], soil communities [23, 24] and the human gut microbiota [25–27].

One critical consideration for field-based metaproteomics studies is the preservation of *in situ* gene expression patterns by stopping biological activity in the samples. In contrast to DNA-based approaches such as 16S rRNA gene sequencing and shotgun metagenomics, for which outcomes only change if cell numbers in the sample change considerably through cell growth and division, protein abundances in cells can change more quickly in response to changing environmental conditions and thus fast preservation in the field is essential for an accurate snapshot of community activity. The time windows in which protein abundance changes are detectable varies greatly between species. For example, for the fast growing *Escherichia coli*, protein abundances have been shown to significantly change within 10-30 minutes after being exposed to environmental stress [28], while changes in slow growing ammonia-oxidizing bacteria take many hours to days to occur [29]. Therefore, sample preservation for metaproteomics should ideally happen within 10 to 30 minutes after removal of a specimen from its environment. While this can easily be achieved in the laboratory, simply by flash freezing the sample, it can be challenging when collecting samples in the field. At remote field sites low-temperature freezers are often unavailable and liquid nitrogen or dry ice for flash freezing are not an option due to transport restrictions to the field site or due to boiling off/sublimation during extended field stays. Therefore, a field compatible method for metaproteomics sample preservation is needed. Such a method should: i) immediately stop biological activity and thus prevent changes in gene expression and protein degradation, ii) preserve the whole metaproteome without bias against specific protein types or taxonomic groups, iii) work for extended storage at room temperature, iv) work for a wide range of sample types and species, v) be non-hazardous and easy to transport, and vi) be available at low cost.

Various studies have evaluated field-compatible preservation methods for nucleic acids in a wide variety of sample types [14, 30, 31]. In contrast, only limited work has been done on field preservation of proteins and, to our knowledge, only for a single cultured bacterial species [32]. Saito et al. examined the performance of SDS extraction buffer, ethanol, trichloroacetic acid (TCA), B-PER and RNA*later***™** to preserve cultures of the marine cyanobacterium *Synechococcus* using flash frozen samples as a control. The test samples were stored at room temperature (RT) for 4 weeks.

The authors found that all samples yielded lower protein concentrations compared to their flash frozen control. Despite this, Saito et al. found that for RNA*later***™** the number of identified proteins and relative protein abundances were highly similar to flash frozen controls while the remaining preservatives showed significantly lower protein identification numbers.

Although studies of different preservation methods and their effects on the protein levels remain limited, the work of Saito et al. [32] suggests that RNA*later***™** is a promising candidate for a field compatible preservative. In fact, RNA*later***™** preservation has been used for (meta)proteomic studies of several bacteria-animal symbioses, however, without proper validation so potential impacts on the metaproteomes remain unknown [33–36].

The objective of our study was to identify and validate a field-compatible preservation method for metaproteomic analyses of bacteria-animal symbioses. We chose to evaluate the protein preservation performances of flash freezing and RNA*later***™** using field-collected samples. To accurately simulate storage of field samples, which are often kept for days to weeks at room temperature due to the lack of refrigeration, we also conducted a time series with RNA*later***™** up to 4 weeks at room temperature. We used the *Olavius algarvensis* symbiosis as a test case. *O. algarvensis* is a gutless marine worm that harbors two aerobic sulfur oxidizing gammaproteobacteria (*Candidatus* Thiosymbion algarvensis (formerly γ1) and γ3-symbiont), two anaerobic sulfate reducing deltaproteobacteria (δ1 and δ4-symbionts) and a spirochaete [37–40]. We chose to use the *O. algarvensis* symbiosis as a model because it has been extensively studied with -omic techniques such as metagenomics, metatranscriptomics and metaproteomics [21, 39, 40]. This provides the benefit of a well-validated test system for which a custom protein sequence database is available for protein identification. Moreover, this symbiosis is highly specific with the animal host always being associated with a set of the same bacterial symbionts, which allowed us to robustly evaluate the preservation performance for both eukaryotic and prokaryotic proteins. Due to its small size (1.5 to 2.5 cm long and 0.1 to 0.13 mm thick) [41], *O. algarvensis* also allowed us to account for samples with little biomass, which is another common limitation when working with field-collected samples. Overall, these data enabled us to provide recommendations to researchers in various fields of biology who work with field-collected samples targeted for metaproteomic analyses.

## Material and Methods

### Sampling and Experimental set up

We conducted two experiments to evaluate the performance of RNA*later***™** for protein preservation of field-collected samples. Throughout this paper we will refer to them as “preservation method comparison” and “RNA*later***™** time series”. In addition, we fully repeated the preservation method comparison to confirm our results from the first experiment.

We collected and processed all samples as previously described [42]. Briefly, samples were collected off the coast of Sant’ Andrea Bay, Elba, Italy (42°48′26″N, 010°08′28″E) from shallow-water (6- to 8-m water depth) sediments next to seagrass beds. We collected 10 individuals of *O. algarvensis* in 2015 for the preservation method comparison, 16 individuals for the replication of the preservation study in August 2019 and 33 individuals for the RNA*later***™** time series in 2016 (Supplemental material Table S1-S3). Live worms were transported in native Elba sediment and seawater to the Max Planck Institute for Marine Microbiology in Bremen, Germany where we carefully removed the worms from the sediment and either froze specimens at −80 °C or immersed them in RNA*later***™** until further processing.

For the preservation method comparison we flash froze five specimens of *O. algarvensis* in liquid nitrogen and then stored them at −80 °C. We incubated the other five individuals in RNA*later***™** (Thermo Fisher Scientific). After 24 hours we removed the RNA*later***™** and stored the samples at −80 °C until processing. We reproduced this set-up in the replication of the preservation method comparison with 9 flash frozen and 7 RNA*later***™** incubated individuals of *O. algarvensis*.

For the RNA*later***™** time series we incubated 33 individuals of *O. algarvensis* in RNA*later***™**. Of these 33 individuals, 11 were incubated for 24 hours in RNA*later***™** at 4°C (t0), while 6, 8 and 8 individuals were incubated in RNA*later***™** at room temperature (22-23 °C) for an additional 24 hours (t1), one week (t2), and four weeks (t3) respectively. We removed RNA*later***™** after incubation and stored the samples at −80 °C until further processing.

### Protein extraction, peptide preparation and determination

Samples for the preservation method comparison, replication of the preservation method comparison and RNA*later***™** time series were processed slightly differently in terms of protein extraction. These steps are described separately for each study. Protein identification and quantification steps were identical in all studies and are thus described only once.

For the preservation method comparison we prepared tryptic peptides following the filter-aided sample preparation (FASP) protocol, adapted from [43]. We added 50 μl of SDT-lysis buffer (4% (w/v) SDS, 100 mM Tris-HCl pH 7.6, 0.1 M DTT) to each sample and heated the samples to 90°C for 10 min. Samples were centrifuged for 5 minutes at 21,000 x g. We mixed 30 μl of each lysate with 200 μl UA solution (8 M urea in 0.1 M Tris/HCl pH 8.5) in a 10 kDa MWCO 500 μl centrifugal filter unit (VWR International) and centrifuged the mixture at 14,000 x g for 40 min. Next, we added 200 μl of UA solution and centrifuged again at 14,000 x g for 40 min. We added 100 μl of IAA solution (0.05 M iodoacetamide in UA solution) and then incubated samples at 22 °C for 20 min in the dark. We removed the IAA solution by centrifugation followed by three wash steps with 100 μl of UA solution. Subsequently, we washed the filters three times with 100 μl of ABC buffer (50 mM ammonium bicarbonate). We added 1.6 μg of Pierce MS grade trypsin (Thermo Fisher Scientific) in 40 μl of ABC buffer to each filter. Filters were incubated overnight in a wet chamber at 37°C. The next day, we eluted the peptides by centrifugation at 14,000 x g for 20 min followed by the addition of 50 μl of 0.5 M NaCl and another centrifugation step. Peptides were quantified using the Pierce MicroBCA Kit (Thermo Fisher Scientific) following the instructions of the manufacturer.

We processed samples of the preservation method replication and the RNA*later***™** time series similar to the preservation method comparison samples with the following modifications: We added 60 μl of SDT-lysis buffer instead of 50 μl and boiled samples at 95°C for 10 min. To minimize sample loss, we did not do the 5 minute centrifugation step at 21,000 x g described in the original protocol [43] and instead mixed the complete 60 μl of each lysate with 400 μl of UA solution in a 10 kDa MWCO 500 μl centrifugal filter unit. All subsequent steps were identical to the sample preparation for the preservation method comparison with the exception that we added 0.62 μg and 0.54 μg of Pierce MS grade trypsin (Thermo Fisher Scientific) in 40 μl of ABC buffer to each filter for the repetition of the preservation method comparison and the RNA*later***™** time series respectively.

### One-dimensional liquid chromatography–tandem mass spectrometry (1D-LC-MS/MS)

All samples were analyzed by 1D-LC-MS/MS. Detailed instrument set-ups, gradients and methods are specified in Additional file S1. In brief, all samples were loaded onto a C18 Acclaim PepMap 100 pre-column and separated on an Easy-Spray PepMap C18, 75μm x 75 cm analytical column (Thermo Fisher Scientific) using reverse phase liquid chromatography. Eluting peptides were ionized with electrospray ionization and mass spectra were acquired using a data dependent acquisition method in a Q-Exactive Orbitrap mass spectrometer (Thermo Fisher Scientific).

### Protein identification and quantification

We downloaded an existing custom protein sequence database for the *O. algarvensis* symbiosis [42] from the PRIDE repository (PXD007510) (http://proteomecentral.proteomexchange.org) and used it for protein identification. The database contained 1,439,794 protein sequences including host and symbiont proteins as well as a cRAP protein sequence database (http://www.thegpm.org/crap/) of common laboratory contaminants. We performed searches of the MS/MS spectra against this database with the Sequest HT node in Proteome Discoverer version 2.2.0.388 (Thermo Fisher Scientific) as described in Gruber-Vodicka et al. [44]. The following parameters were used: trypsin (full), maximum two missed cleavages, 10 ppm precursor mass tolerance, 0.1 Da fragment mass tolerance and maximum of 3 equal dynamic modifications per peptide, namely: oxidation on M (+ 15.995 Da), carbamidomethyl on C (+ 57.021 Da) and acetyl on the protein N terminus (+ 42.011 Da). False discovery rates (FDRs) for peptide spectral matches (PSMs) were calculated and filtered using the Percolator Node in Proteome Discoverer [45]. Percolator was run with a maximum delta Cn 0.05, a strict target FDR of 0.01, a relaxed target FDR of 0.05 and validation based on q-value. The Protein FDR Validator Node in Proteome Discoverer was used to calculate q-values for inferred proteins based on the results from a search against a target-decoy database. Proteins with a q-value of <0.01 were categorized as high-confidence identifications and proteins with a q-value of 0.01–0.05 were categorized as medium-confidence identifications. We combined search results for all samples into a multiconsensus report in Proteome Discoverer and only proteins identified with medium or high confidence were retained, resulting in an overall protein-level FDR of 5%. For protein quantification, normalized spectral abundance factors (NSAFs, [46]) were calculated per species and multiplied by 100, to give the relative protein abundance in %.

### Outlier identification and removal

We classified samples as outlier if at least two out of the following criteria were met i) the total ion chromatogram intensity was below 1×10^9^; ii) the proportional number of standard deviations above and below the mean (z-score) of the number of identified proteins (filtered for 5% FDR) was > ∓ 1; iii) the number of identified proteins (filtered for 5% FDR) was more than one standard deviation below the mean number of identified proteins of all samples within a group (Additional file S2). In addition, we also applied the Generalized Extreme Studentized deviate test (ESD) (significance level of 0.5, maximum of 10 outliers) on the number identified proteins in the RNA*later***™** time series for outlier identification. This procedure was not applied on the preservation method comparison and the replication of the preservation method comparison due to insufficient number of replicates. In total, we identified 2 samples of the preservation methods comparison, 2 samples of the repetition of the methods comparisons and 8 samples of the RNA*later***™** time series as outliers (Additional file S2). Identified outliers were excluded from all subsequent analyses.

In addition, we checked the metaproteomes for evidence of accidental sampling of the co-occurring marine gutless oligochaete *Olavius ilvae* [47]. *O. ilvae* cannot be easily distinguished from *O. algarvensis* during sampling as *O. algarvensis* and *O. ilvae* are highly similar in size, shape and color. However, they harbor distinct symbionts, which can be used to distinguish between the species [48]. To test whether any of our samples was a specimen of *O. ilvae*, we created a custom database including protein sequences of the α7-symbiont, *Cand*. Thiosymbion sp., γ3-symbiont and δ3-symbiont of *O. ilvae*. In addition, we also included protein sequences of *Cand*. Thiosymbion algarvensis and the δ1-symbiont of *O. algarvensis* for testing. The database was then loaded into Proteome discoverer and proteins were identified as described above. One sample of the RNA*later***™** time series was identified as *O. ilvae* and therefore removed as an outlier (Additional file S2).

### Data analysis

To determine which identified proteins were shared by all samples or unique to specific treatments/time points we loaded the 5% FDR filtered PSM multiconsensus files into Perseus 1.6.5.0 (Tyanova et al., 2016), filtered out proteins that did not have at least 75% valid values (greater than 0) in at least one group and log2 transformed the data. We then calculated the overlap protein sets with the numerical Venn function in Perseus and visualized the results with a Venn calculation tool from Ghent University (http://bioinformatics.psb.ugent.be/webtools/Venn/) using the default settings.

For hierarchical clustering we loaded the 5% FDR filtered NSAF multiconsensus files into Perseus and filtered out proteins that did not have at least 75% valid values in at least one group and log2 transformed the data. We replaced invalid values with a constant value and z-score normalized the resulting matrix by rows (proteins). Subsequently, we performed hierarchical clustering with the following settings: Euclidean distance, preprocessing with k-means and average linkage.

For differential protein abundances we loaded the 5% FDR filtered NSAF multiconsensus files in Perseus. We grouped samples by preservation method/ time point and filtered proteins for 75% valid values in at least one group to only use consistently identified proteins. We replaced missing values by a constant and performed a two-sided Welch’s t-test using a permutation-based false discovery rate of 5% to account for multiple hypothesis testing.

For the differential protein abundance analysis of the 1000 most abundant proteins we loaded the 5% filtered NSAF multiconsensus, calculated the NSAF sum across all treatments and sorted proteins from most to least abundant. We selected the 1000 most abundant proteins and re-normalized the abundance values based on the selected subset. We then loaded the resulting matrix in Perseus, grouped samples by preservation method/ time point, log2 transformed the data and replaced missing values by a constant. We used the resulting matrix as input data for Volcano plots based on a t-test with a FDR of 0.05 and S0 of 0.1.

We calculated relative abundances for each species in the symbiosis using the method for assessing the proteinaceous biomass described by Kleiner et al. [49], with the following modification. Instead of using FidoCT for protein inference in Proteome Discoverer and filtering for proteins with at least 2 protein unique peptides, we used Sequest HT for protein inference and filtered for proteins with at least 2 protein unique peptides. We visualized the results with the ggplot package in R [50, 51].

To assess biochemical properties of all identified proteins we obtained the number of amino acids and predicted isoelectric points (pI) from Proteome Discoverer. We predicted transmembrane helices (TMHs) with the TMHMM Server 2.0 [52]. Protein sequences of all identified proteins for each study were used as input data.

### Data availability

The metaproteomics mass spectrometry data and protein sequence database have been deposited to the ProteomeXchange Consortium via the PRIDE partner repository [53] with the following dataset identifiers: Preservation method comparison PXD014591 (Reviewer access at: https://www.ebi.ac.uk/pride/login: Username, Password: qSFUOntG; Replication of the method preservation method comparison PXD026631 (Reviewer access at https://www.ebi.ac.uk/pride/login: Username, Password: OZu9Zs9K), RNA*later***™** time series PXD014881 (Review access at https://www.ebi.ac.uk/pride/login Username, Password: d5MenBMF; *E. coli* grown under oxic and anoxic conditions PXD024288 (Reviewer access at: https://www.ebi.ac.uk/pride/login Username: Password: oQcd2WjH).

## Results

We compared flash freezing and RNA*later***™** preservation to determine if RNA*later***™** is a suitable method for preservation of field-collected samples targeted for metaproteomic analyses. We used the marine gutless oligochaete *O. algarvensis* and its bacterial endosymbionts as our test system. To simulate field conditions, we also conducted a time series to assess how metaproteomes were affected by storage of samples in RNA*later***™** for up to 4 weeks at room temperature.

### Similar numbers of proteins identified for both preservation methods and all RNA*later*™ storage time points

We identified similar numbers of proteins for both flash frozen and RNA*later***™** preserved samples. On average, we identified 5,934 proteins in flash frozen samples and 5,780 proteins in RNA*later***™** preserved samples (Fig. 1A). The average number of identified proteins between flash frozen samples and RNA*later***™** preserved samples was not significantly different (student’s t-test, p > 0.05). This suggests that neither of the tested methods outperforms the other in terms of total number of identified proteins.

**Fig. 1:**
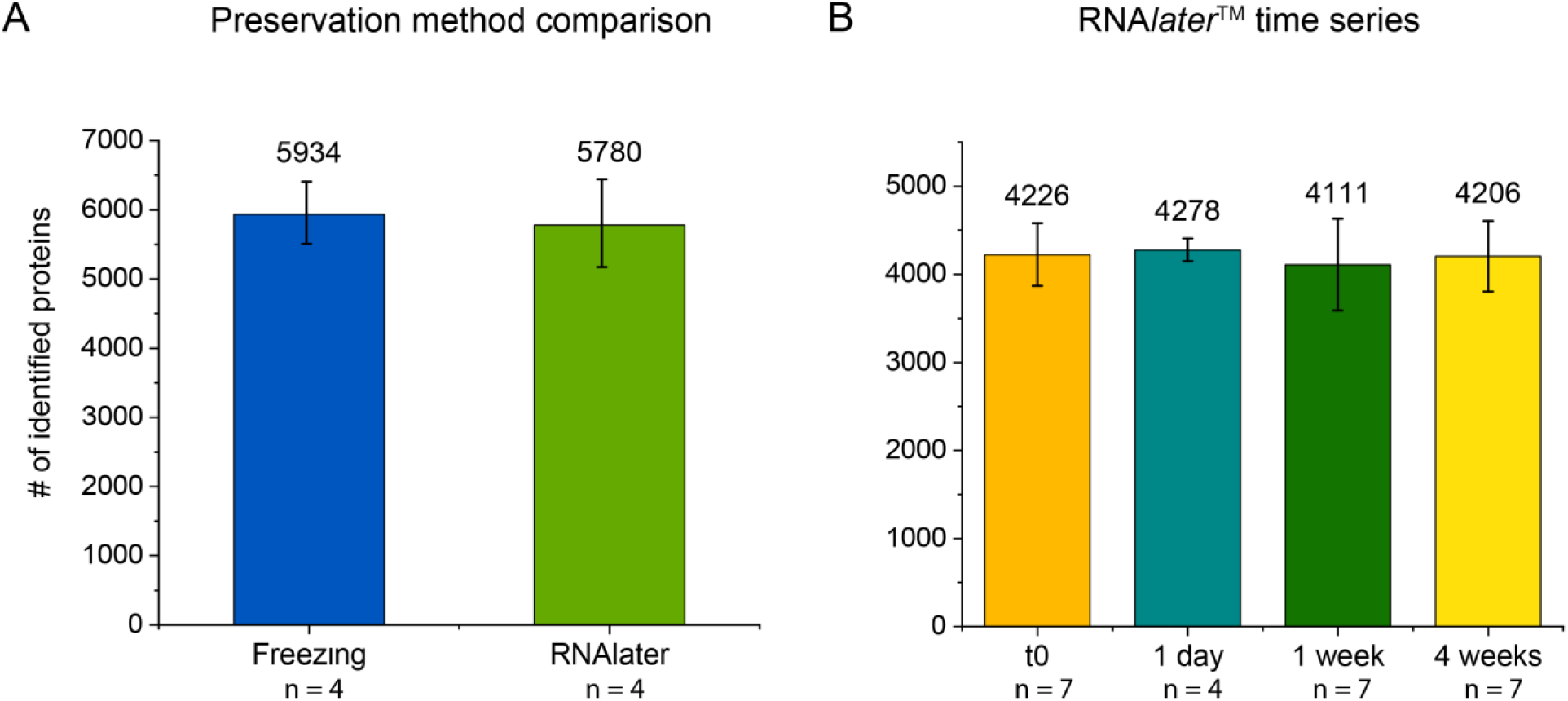
Number of identified proteins did not differ between preservation methods and time points. Identified proteins were filtered for a false discovery rate (FDR) of 5% prior to counting. Error bars indicate the standard deviation of the mean (SD). **A)** Average number of identified proteins from flash frozen and RNA*later***™** preserved samples. **B)** Average number of identified proteins in samples from the four RNA*later***™** storage time points. Samples were frozen immediately after incubation in RNA*later***™** at 4°C overnight for the t0 time point and after 1 extra day, 1 week and 4 weeks of RNA*later***™** incubation at room temperature. It is noteworthy to mention that the number of total identified proteins between A) and B) varies due to small but relevant differences in the protein extraction protocols. These differences led to more efficient extraction of abundant muscle proteins of the host in samples of the RNA*later***™** time series. The high overabundance of peptides from these muscle proteins increased the detection limit for other, lower abundant proteins.

The number of identified proteins was stable across the four tested storage time points. On average, we identified between 4,111 and 4,278 proteins per time point (Fig. 1B). None of the total protein numbers were significantly different as compared to t0, the starting point of the RNA*later*™ incubation (student’s t-test, p > 0.05). While the manufacturer recommends that samples should be stored at 4°C if storage exceeds 1 week our results suggest that proteins are well preserved for at least 4 weeks at room temperature.

### No systematic bias in protein identification and quantification between preservation methods and storage time points

Our results show that there is no systematic bias when sample preservation methods and time points are compared based on overall protein identification and quantification. To identify systematic bias we analyzed the data for proteins consistently present in only one of the preservation methods or time points, and also applied hierarchical clustering based on protein abundances to check if samples formed groups based on method or time point (Fig. 2 A-D). For these analyses we only included proteins that were consistently detected in at least one of the treatments/time points by filtering out proteins that were not detected in at least 75% of samples for at least one condition.

**Fig. 2:**
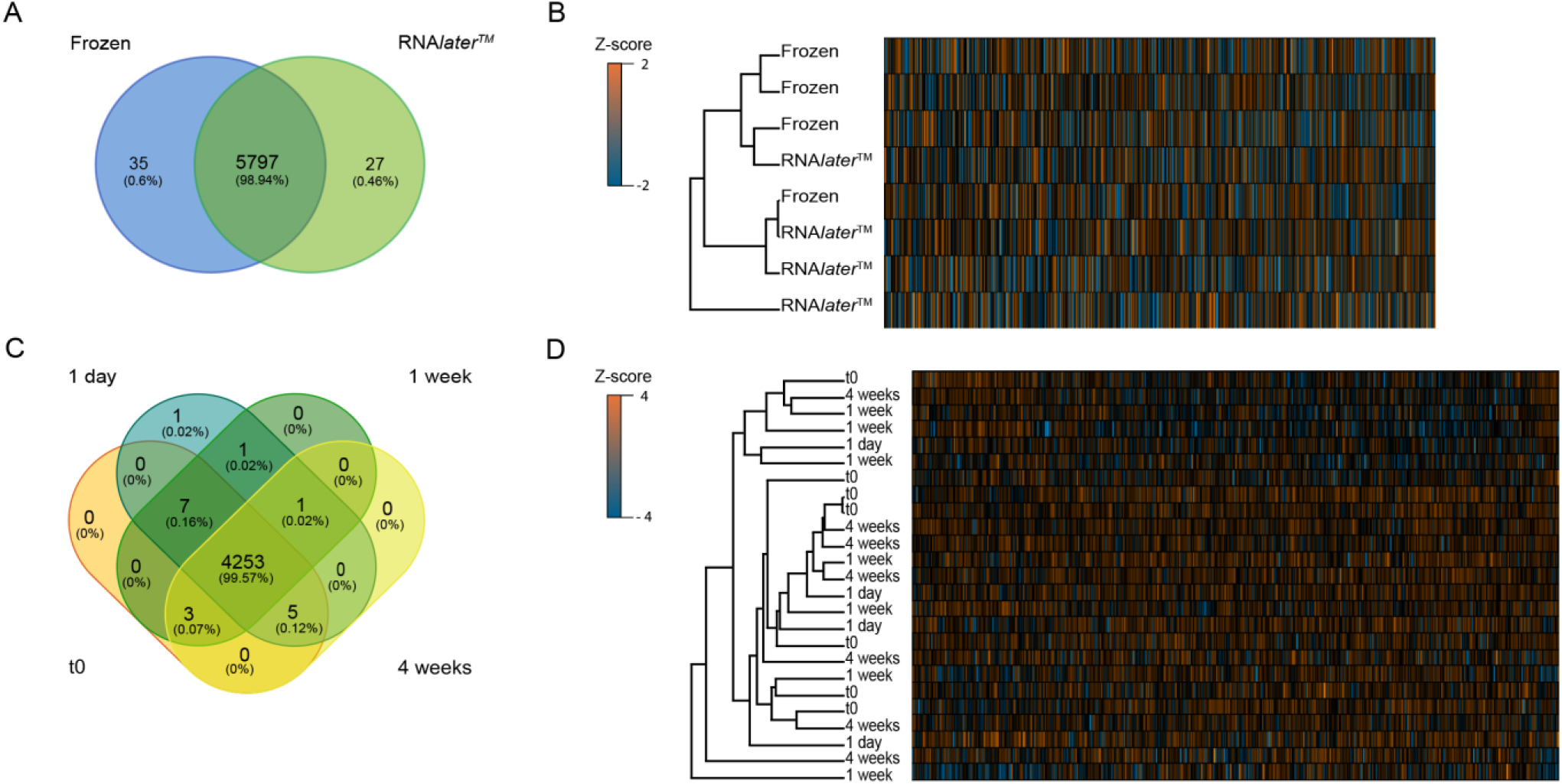
No systematic bias in protein identification and quantification between preservation methods and storage time points. Unique and shared proteins identified in the preservation method comparison **A)** and the RNA*later***™** time series **C).** Only proteins that were retained after filtering for a 5% FDR and that were detected in at least 75% of samples in at least one group (preservation method/time point) were used for the analysis. Hierarchical clustering of samples based on relative protein abundances for the preservation method comparison **B)** and the RNA*later***™** time series **D)** The same set of pre-filtered proteins was used as in A and C and log2 transformed. The hierarchical clustering was based on Euclidean distance. Z-score values were calculated for each protein and thus positive Z-scores indicate a relative abundance higher than the mean, while negative Z-scores indicate a relative abundance lower than the mean.

Of 9,326 proteins identified in the preservation methods comparison (Additional file S3), 5,859 proteins remained after filtering and thus were considered to be consistently identified in at least one of the treatments. Out of these 5,859 proteins almost all (5,797) were shared between flash frozen samples and samples preserved in RNA*later***™** (Fig. 2A). The hierarchical clustering of these samples based on protein abundances revealed multiple shared nodes in the dendrogram between flash frozen and RNA*later***™** preserved samples (Fig. 2B). In case of a systematic bias introduced by the preservation method we would expect separation of samples based on preservation method with no shared nodes between preservation methods. These data suggest that the preservation methods did not introduce a systematic bias.

Of 7,036 proteins identified in the RNA*later***™** time series (Additional file S4), 4,271 were consistently identified and used for the overlap analysis. Out of these 4,271 proteins, 4,253 proteins were shared across all four storage time points (Fig. 2 C). We identified 1 unique protein for samples incubated for 1 day, whereas none of the other time points had unique proteins. Moreover, a few proteins were shared between two or three out of the four different time points. The hierarchical clustering of these samples revealed multiple shared nodes between samples of all time points (1 day, 1 week and 4 weeks) and samples of t0 (Fig. 2D). If a systematic bias had been introduced by long-term storage at room temperature we would expect separation of samples based on time point with no shared nodes between test time points and t0. These data suggest that long-term storage at room temperature did not introduce a systematic bias.

### Only minor differences detected in relative abundances of individual proteins across preservation methods or time points

We evaluated relative protein abundances across preservation methods and storage time points to assess potential alterations in protein abundances introduced by method or time. For these analyses we used the same dataset as above, including only proteins that were consistently detected in at least one of the treatments/time points. We used a two–sided Welch test to identify significant differences in protein abundances between methods and time points.

Out of the 5,859 consistently identified proteins in the preservation method comparison, 14 proteins had abundances that significantly differed between treatments. Out of these 14 proteins, 13 proteins were host proteins and 1 protein was a δ1-symbiont protein (Additional file S5). To investigate whether the 14 proteins shared any characteristics that might explain their differing abundances, we checked their overall abundances (ranking positions), protein lengths, isoelectric points (pI) and numbers of predicted transmembrane helices (TMHs) (Additional file S5). No clear trend could be observed other than the host proteins had relatively low abundances.

Out of the 4,789 consistently identified proteins in the RNA*later***™** time series, 2 proteins were significantly different between samples incubated for 1 day in RNA*later***™** compared to t0. These two proteins were both host proteins with no clear pattern in regards to their biochemical properties. No protein abundances significantly differed between samples incubated for 1 week and 4 weeks.

We selected the 1000 most abundant proteins from each study and calculated the average relative protein abundances for each method or time point to lower the influence of biological variability between individual worms. We log2 transformed the data and visualized the differences between with Volcano plots based on a t-test with a FDR of 0.05 and S0 of 0.1 (Fig. 3 A-E). The S0 parameter indicates the relative importance of t-test p-value meaning that even if a protein has a significant p-value, if the fold change is below the S0 value it will not be considered statistically significant. If proteins abundances significantly differed between treatments/time points they would appear above the S0 line in the plot whereas if their abundances did not significantly differ their values would be below the S0 line. We were unable to identify any significant differences in the 1000 most abundant proteins identified in the preservation method comparison (Fig 3A) or RNA*later***™** time series (Fig. 3B-D). To further emphasize this finding we included an example study of *Escherichia coli* which was grown in either oxic or anoxic conditions (Fig. 3E). As expected, there were several differentially expressed proteins between the two growth conditions as indicated by data points above the S0 line in Figure 3E. In summary, our analysis revealed no significant changes in protein abundances between the tested preservation methods, as well as between t0 and the storage time points of the RNA*later***™** time series.

**Fig. 3:**
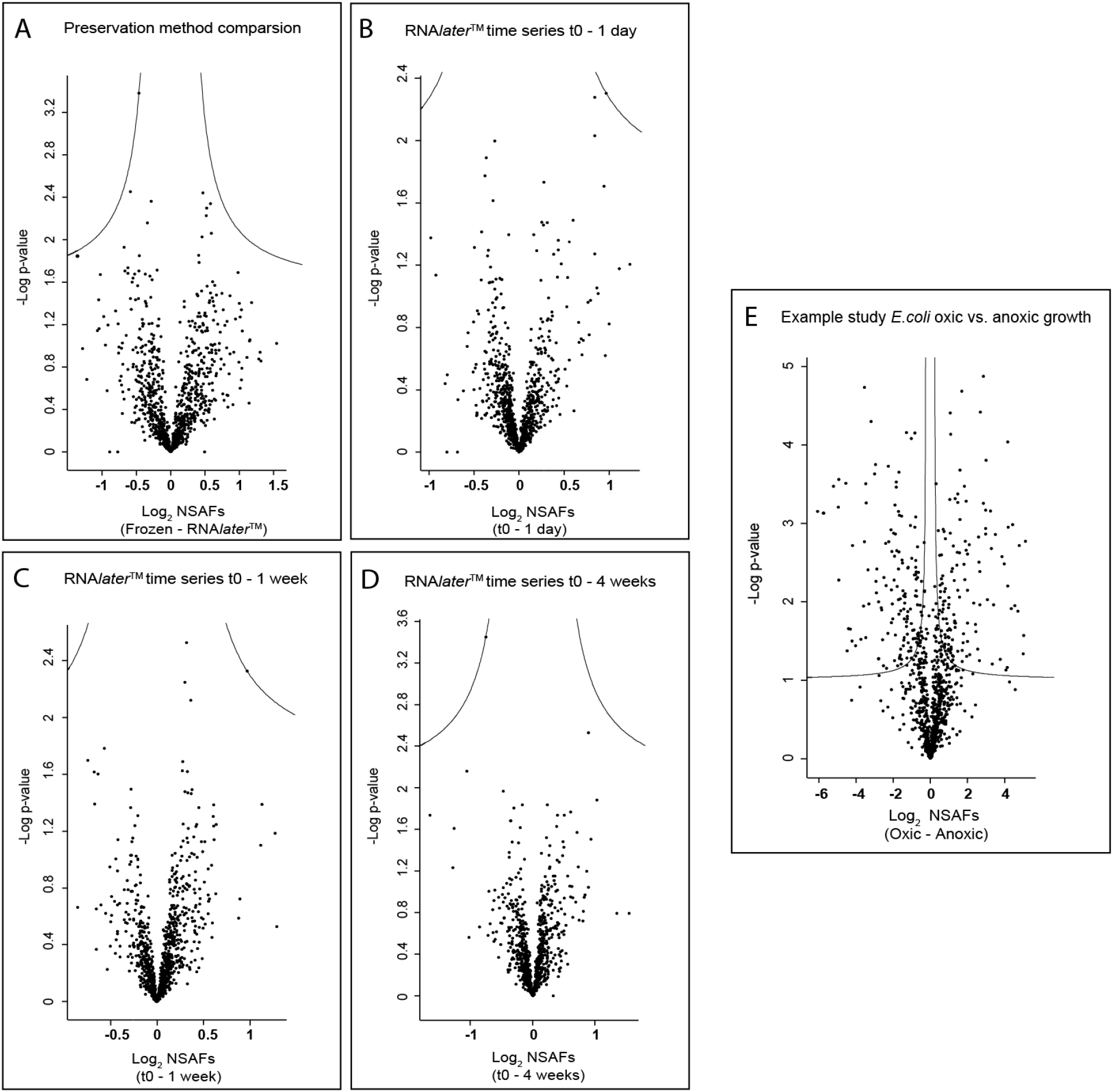
No significant differences in protein abundances between preservation methods and storage time points for the 1000 most abundant proteins. Volcano plots of the 1000 most abundant proteins identified in the preservation method comparison **A)**, the RNA*later***™** time series **B-D)** and an example study of *E.coli* grown in oxic and anoxic conditions **E)**. All proteins were plotted with log 2 fold change on the x-axis and – log (p- value) on the y-axis. The solid lines represent S0 with a value of 0.1. Data points above the line represent proteins whose abundances significantly differed between comparisons whereas data points below the line represent proteins whose abundances did not significantly differ between comparisons.

### Effects on microbial community structure

To investigate potential effects of preservation method or storage time on the representation of specific taxa in the metaproteome, we compared the proteinaceous biomass of each community member using a method adapted from Kleiner et al. [49]. This method enables calculations of proteinaceous biomass contributions of species in microbial communities by using protein abundances derived from metaproteomic analyses.

We found a small but significant difference in proteinaceous biomass of the host and *Cand*. T. algarvensis in the preservation methods comparison (student’s t-test, p-value < 0.05) (Fig. 4A, Supplemental material Table S4-5). In flash frozen samples the host’s biomass accounted for an average of 75.42% of total biomass while it was an average of 82.40% in RNA*later***™** preserved samples. In contrast to the host’s biomass, the average biomass of *Cand*. T. algarvensis was higher in flash frozen samples (17.61%) compared to RNA*later***™** preserved specimens (13.48%). None of the other symbionts showed a significant difference. To confirm if preservation method truly affected representation of specific taxa we repeated this experiment with a fresh set of 16 *O. algarvensis* individuals to exclude biological variability between individual worms as a potential reason for the observed difference (Fig. 4B, Supplemental material Table S3, Supplemental material Figure S1). In this repeat experiment of the preservation method comparison we did not observe any significant differences in proteinaceous biomass for the host or *Cand*. T. algarvensis or any other taxa (Fig.5 B, Supplemental material Table S6-7, Figure S1). This suggests that there is either no effect or a small inconsistently occurring effect on taxa representation introduced by flash freezing and RNA*later***™** preservation.

**Fig. 4:**
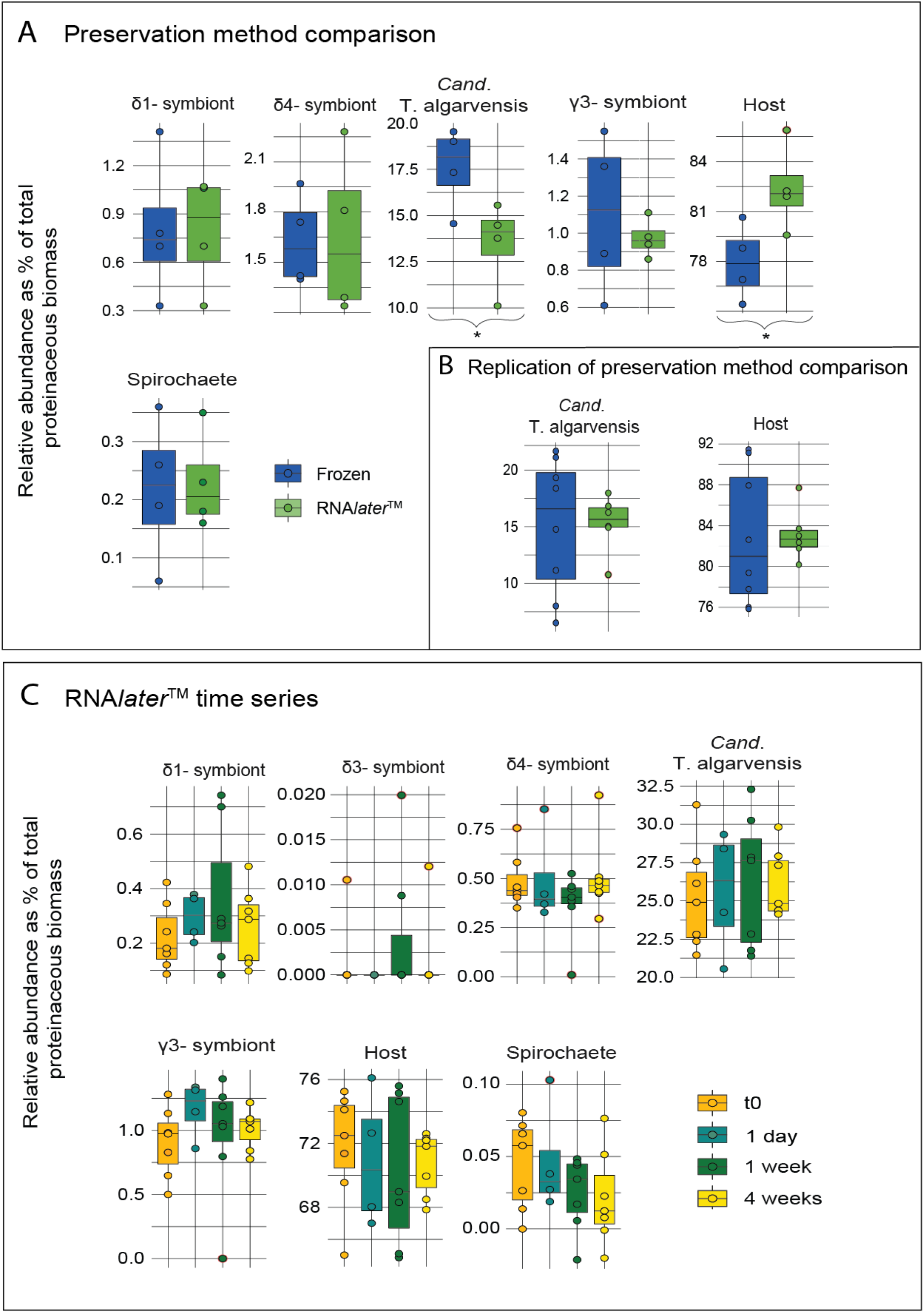
Per-species biomass estimates for members of the *O. algarvensis* symbiosis are mostly consistent for preservation methods and RNA*later*™ time series time points. Data for individual *O. algarvensis* specimens are shown for the preservation method comparison **A)**, a subset of the repetition of the preservation method comparison **B)** and all storage time points of the RNA*later***™** time series **c)**. Taxa were quantified using the sum of PSM counts for each species respectively. Identified proteins were filtered for 5% FDR and at least 2 protein unique peptides (PUP) prior to counting as described in [49]. Asterisks indicate significant differences in per-species biomass (student’s t-test, p-value < 0.05) (Supplemental material Table S4-9).

**Fig. 5:**
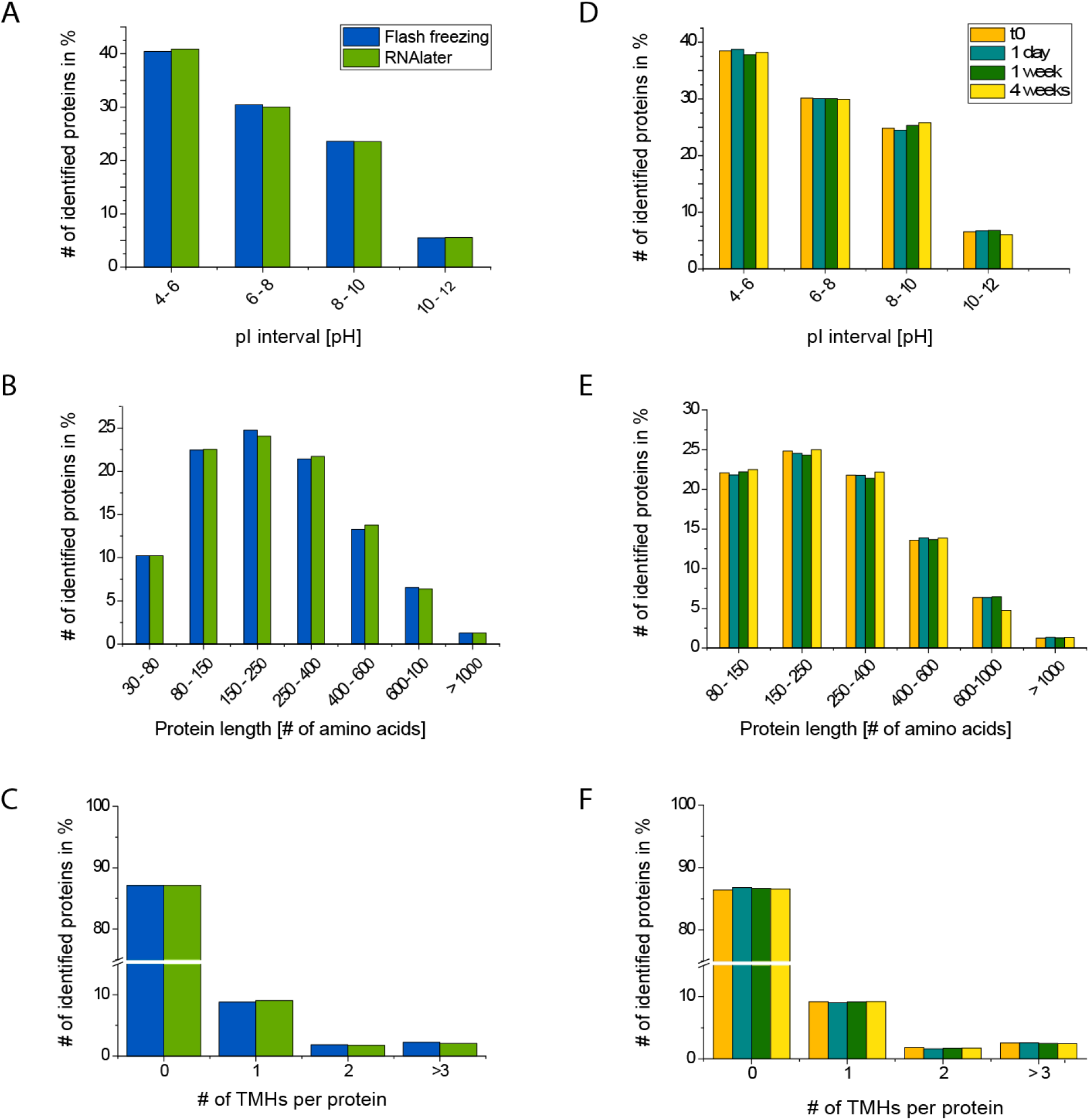
Preservation method and storage time in RNA*later*™ does not introduce any biases against proteins with specific biochemical properties. Identified proteins were filtered for 5% FDR prior to counting. Only intervals contributing more than 1% of all identified proteins are displayed (e.g. pI 2-4 is not shown). Isoelectric point (pI) distribution of identified proteins for the preservation method comparison **A)** and the RNA*later***™** time series **D)**. Protein length distribution for the preservation method comparison **B)** and the RNA*later***™** time series **E)** Number of predicted transmembrane helices (TMHs) for the preservation method comparison **C)** and the RNA*later***™** time series **F)**.

In the RNA*later***™** time series, measured biomass abundances of species were relatively consistent between individual worms (Fig. 4C, Supplemental material Table S8). The only exception was the δ3-symbiont that was detected in only 3 individuals, which is in line with the fact that this symbiont has been shown to be only present in a minority of individuals [8]. None of the symbiont or host biomasses were significantly different when later time points were compared to t0 (student’s t-test, p-value < 0.05) (Fig. 4C, Supplemental material Table S9) indicating that storage up to 4 weeks in RNA*later***™** did not impact proteinaceous biomass measurements of individual species.

### No evidence for biases against proteins with specific biochemical properties

The two tested preservation methods rely on distinct preservation mechanisms, which holds the potential for categorical loss or enrichment of proteins based on their biochemical properties. To evaluate the potential introduction of method or storage time specific biases, we evaluated biochemical properties including protein size, isoelectric point (pI) and number of transmembrane helix domains (TMHs) across all samples. In contrast to the overlap analysis shown in Figure 2 A and C, for this analysis all proteins identified within an FDR of 5% were considered.

We did not observe any significant differences in pI, protein size or number of predicted transmembrane helices (TMHs) for preservation methods (Fig. 6A-C, student’s t-test, p-value < 0.05) and storage time points (Fig. 6 D-F, student’s t-test, p-value < 0.05). Counts within the respective intervals were almost identical for all examined parameters. For example, the average pI of proteins in flash frozen samples was 6.86 while it was 6.85 in RNA*later***™** preserved samples and the mean protein length was 296 amino acids for frozen samples and 297 amino acids for RNA*later***™** preserved samples. Overall, our analysis showed that we recovered proteins with almost identical biochemical properties for both preservation methods and all storage time points. This suggests that RNA*later***™** robustly preserves proteins at room temperature for at least 4 weeks without introducing biases based on biochemical properties.

## Discussion

We evaluated the protein preservation performance of flash freezing in liquid nitrogen and RNA*later***™** for metaproteomics of field-collected samples. Our main finding was that both preservation methods performed equally well and that storage time in RNA*later***™** for up to 4 weeks did not impact the quality of the metaproteomes. However, there were other parameters which need to be taken into account when considering actual field deployment, some of which might vary depending on the sampling location and experimental design (Table 1).

**Table 1:**
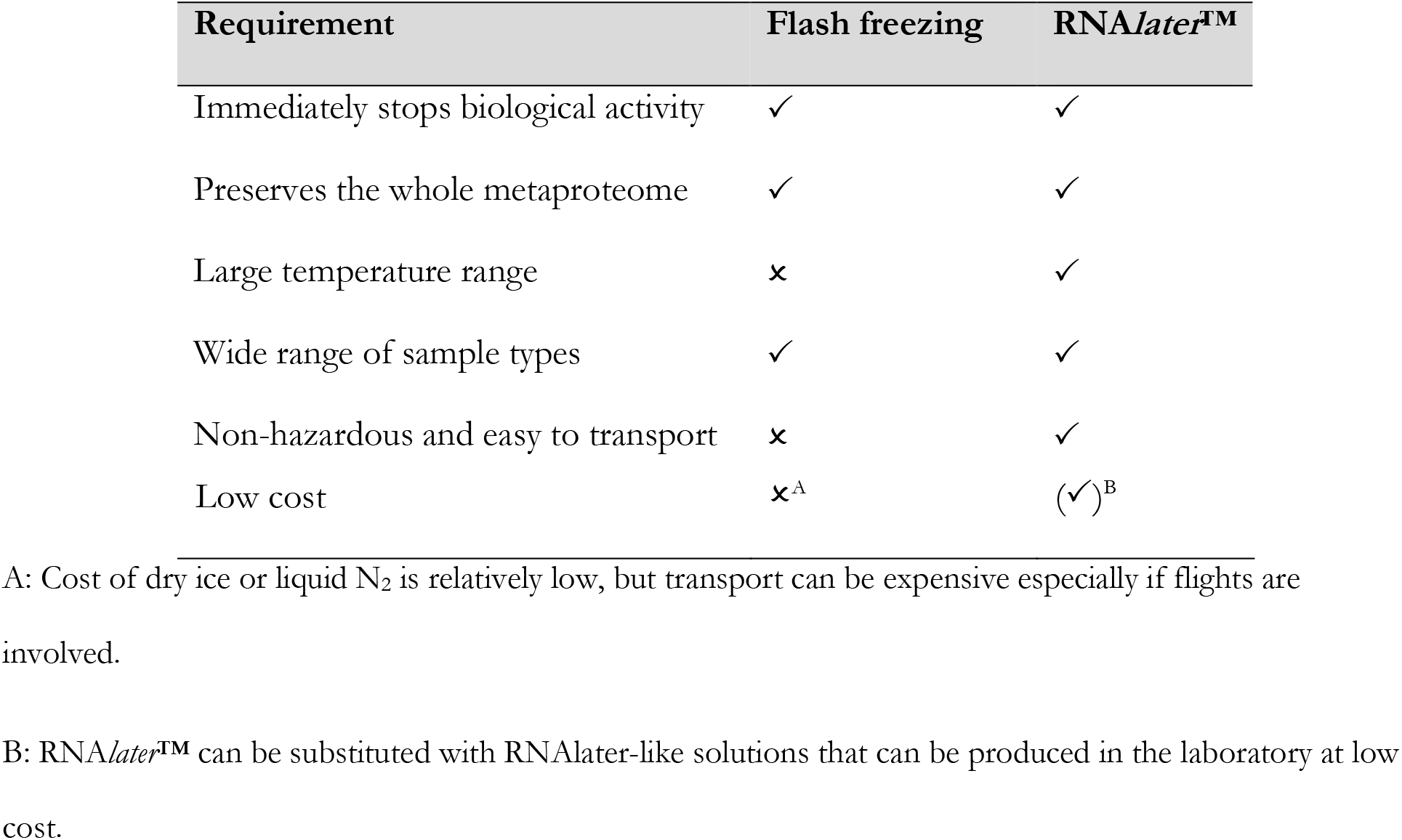
Comparison of suitability of flash freezing in liquid nitrogen and RNA*later***™** for protein preservation in field deployment.

The comparison of flash freezing and preservation in RNA*later***™** showed that both methods stop biological activity equally well. In addition, we found no alteration in preservation performance of RNA*later***™** over time and at room temperature. Flash freezing inhibits biological activity through a fast freezing process at ultra-low temperatures (−195.79 °C) while RNA*later***™** contains high concentrations of ammonium sulfate which causes protein precipitation and hence inactivation. Depending on the sample type and size, RNA*later***™** might take a few minutes to completely immerse the sample. While one could argue that this time holds the potential to introduce biases and protein degradation, our analyses showed that this concern is unsubstantiated. However, we recommend these results be re-validated when working with larger animals or animals with an impermeable exoskeleton as dissection into smaller pieces than the manufacturer’s recommendation 0.5 cm may be required for a fast and sufficient infusion of tissue.

Considering actual field deployment, an important parameter for a protein preservative is its ability to work at large temperature ranges as weather conditions might vary significantly depending on the sampling location. We found that samples immersed in RNA*later***™** were robustly preserved up to 4 weeks at room temperature (22-23 °C). Generally, when RNA*later***™** is used for RNA preservation it has been shown to perform better when at lower temperature (e.g. 4 °C) [54]. We thus recommend storing samples as near this temperature as possible, and consider 23 °C the upper temperature limit for longer-term sample storage. If field conditions require longer-term storage above 23 °C we recommend re-testing of the impact of storage on metaproteome quality.

Another important aspect for field-compatible protein preservatives is that they should be non-hazardous and easy to transport to remote field locations. While this is the case for RNA*later***™**, which can be stored in plastic containers at room temperature, this does not hold true for liquid nitrogen which can cause cryogenic burns, injury or frostbite and may displace oxygen and cause rapid suffocation if handled improperly. As a consequence, a liquid nitrogen shipment requires thorough planning prior to the sampling trip to ensure a safe and smooth arrival at the destination. We were actually inspired to do this study in part because, in the past, despite thorough planning, our liquid nitrogen dry shippers did not arrive at the field site several times. Moreover, a cryogenic shipment can easily exceed reasonable costs since it requires a field compatible cryogenic dry shipper and comes with high shipping cost, although the cost of liquid nitrogen itself is relatively low ($0.50-2.00 per liter). RNA*later***™**, on the other hand, is comparably expensive ($477 for 500 mL, Thermo Fisher Website on 21 Feb 2021); however, the amount of RNA*later***™** needed per sample is potentially very small as the required RNA*later***™** to sample ratio is 10:1. This means, for example, for the very small *O. algarvensis* worms less than 100 μl of RNA*later***™** were needed per sample. Additionally, the high cost of RNA*later***™** can be avoided by using a self-prepared RNAlater-like solution which has been shown to perform as well as the commercially available solution [55].

Finally, once samples are brought back to the laboratory to await processing, they should either be stored at 4°C within RNA*later***™**, or alternatively, if samples are targeted for long-term storage, RNA*later***™** should be removed and samples should be frozen at −80 °C. When removing the RNA*later***™** prior to freezing the samples, do not wash the samples with a buffer as the RNA*later***™** precipitation effect is reversible and proteases would become active again upon dilution of RNA*later***™**.

## Ethics approval and consent to participate

Not applicable.

## Availability of data and material

The mass spectrometry metaproteomics data and protein sequence databases have been deposited to the ProteomeXchange Consortium via the PRIDE partner repository. Data set identifiers are PXD014591, PXD026631 and PXD014881 for the preservation method comparison, the replication of the preservation method comparison and the RNA*later***™** time series respectively.

## Authors’ contributions

M.J., M.K. and J.W. designed the study. M.J., J.W. and M.K. performed experiments. M.J performed data analyses. M.J. and M.K. wrote the manuscript. All authors read and approved the final manuscript.

## Acknowledgements

We thank Marc Strous for access to proteomics equipment. Part of the LC-MS/MS measurements were made in the Molecular Education, Technology, and Research Innovation Center (METRIC) at North Carolina State University. We also thank Angie Mordant and Fernanda Salvato for help with protein extraction and technical assistance, Nicole Dubilier for provision of supplies and access to her laboratory at the Max-Planck-Institute for Marine Microbiology, and Dolma Michellod and Tina Enders for providing help with sample collection.

